# Deciphering the Gut–Brain Dialogue: A Survey-Based and *In-Silico* Comparative Analysis of Gut Microbial Dysbiosis in Common Neurological Disorders

**DOI:** 10.64898/2026.08.24.746718

**Authors:** Shreya Goyal, Aakanksha Kalra

## Abstract

The gut microbiome maintains a complex, bidirectional communication network with the central nervous system, commonly referred to as the gut-brain axis and its disruption has been implicated in several neurological disorders. This study combines a survey-based assessment of public awareness with an *in-silico* comparative analysis of gut microbial dysbiosis across four prevalent neurological disorders as observed in the current study: depression, anxiety, schizophrenia and autism spectrum disorder (ASD). A structured, anonymous online survey (n = 230) captured perceptions of the gut-brain connection along with dietary, lifestyle and gastrointestinal correlates of stress in a predominantly young, health-sciences-affiliated Indian cohort. In parallel, disorder-specific lists of elevated and reduced faecal microbial taxa were retrieved from the Disbiome database, compared using a multiple list comparator and taxonomically classified using the NCBI Taxonomy tool to construct phylogenetic trees in iTOL. Approximately three-quarters of respondents were aware of a potential gut-mental health link, yet about half reported no specific dietary practice and roughly 60% experienced stress-related digestive symptoms while rarely seeking medical consultation for them. Comparative analysis showed that depression, anxiety and schizophrenia shared a substantially overlapping dysbiosis signature, with common elevation of Actinomyces, Bacteroidaceae, Blautia, Eggerthella, Oscillibacter, Parasutterella and Veillonella and common reduction of Coprococcus, Lachnospiraceae, Ruminococcaceae, Clostridium, Faecalibacterium and Sutterella. In contrast, ASD displayed a distinct microbial signature with limited overlap with the other three disorders. Phylogenetic clustering confirmed that the shared taxa belonged predominantly to the phyla *Bacillota* (formerly *Firmicutes*), *Bacteroidota* (formerly *Bacteroidetes*), *Actinomycetota* (formerly *Actinobacteria*) and *Pseudomonadota* (formerly *Proteobacteria*). Notably, this phylum-level pattern parallels recent comparative analyses of microbial dysbiosis in neurodegenerative diseases, suggesting that broad phylogenetic shifts may be a relatively general correlate of chronic neurological disease, while disorder specificity emerges at the level of individual taxa. These findings support a shared microbial pathway linking depression, anxiety and schizophrenia that is distinct from the dysbiosis pattern observed in ASD and they underscore the value of microbiome-informed, disorder-specific therapeutic strategies.

## 1. Introduction

The human gastrointestinal tract harbours trillions of bacteria, archaea, fungi and viruses that collectively constitute the gut microbiome. Beyond its established roles in digestion, nutrient absorption and vitamin synthesis, this microbial community actively shapes host immunity and provides a barrier against pathogenic colonization [1]. A stable, diverse microbiota is a hallmark of health, whereas disease states are frequently accompanied by a shift toward pathogenic taxa, a condition termed microbial dysbiosis, which has been linked to metabolic, inflammatory and neurological disorders [1].

Mental and neurological disorders constitute a substantial share of the global disease burden. Depression and anxiety alone affect more than 260 million and 280 million people worldwide, respectively and India’s National Mental Health Survey (2015–16) estimated that nearly 15% of Indian adults experience some form of mental illness [2,4,6]. Autism spectrum disorder (ASD) and schizophrenia, though less prevalent, are associated with considerable lifelong disability, affecting approximately 1 in 100 children and 0.3-0.7% of the population in India, respectively [4,6]. Given the scale of this burden, mechanistic insight that could inform new diagnostic or therapeutic avenues is urgently needed.

Mental and neurological disorders constitute a substantial share of the global disease burden. Depression and anxiety alone affect more than 260 million and 280 million people worldwide, respectively, and India’s National Mental Health Survey (2015–16) estimated that nearly 15% of Indian adults experience some form of mental illness [2,4,6]. Autism spectrum disorder (ASD) and schizophrenia, though less prevalent, are associated with considerable lifelong disability, affecting approximately 1 in 100 children and 0.3–0.7% of the population in India, respectively [4,6]. These four disorders were chosen because they span distinct points on the gut–brain axis spectrum: depression and anxiety are prevalent, frequently co-occurring mood disorders; schizophrenia is a severe, later-onset psychotic disorder; and ASD is a neurodevelopmental condition emerging while the gut microbiome itself is still being established. Each has independently been linked to gut dysbiosis, with sufficient curated data in the Disbiome database to allow systematic comparison. Given this burden, and the largely unexplored question of whether these disorders share a common microbial signature, mechanistic insight that could inform new diagnostic or therapeutic strategies is urgently needed.

A growing body of evidence implicates the gut-brain axis, a bidirectional communication network linking the gastrointestinal tract and the central nervous system through neural (principally vagal), endocrine and immune pathways, in the pathophysiology of several neuropsychiatric conditions [3,7]. Gut bacteria synthesise and modulate neurotransmitters, including serotonin, dopamine and γ-aminobutyric acid (GABA), regulate the hypothalamic-pituitary-adrenal (HPA) stress axis and influence systemic and neuro-inflammation through microbial metabolites such as short-chain fatty acids (SCFAs) and pro-inflammatory cytokines [3,7,8]. Disruption of this axis in early life can also compromise blood-brain barrier integrity and synaptic plasticity, with lasting consequences for neurodevelopment [9]. Distinct, disorder-specific patterns of dysbiosis have been reported for depression, anxiety, ASD and schizophrenia individually [10,11,12,13], yet whether these disorders share a common microbial signature, or instead reflect largely independent perturbations, has rarely been examined through direct, cross-disorder comparison.

This study is therefore conducted via a two-part investigation. First, an online survey was conducted to gauge current public understanding of the relationship between gut health and mental well-being, together with associated dietary, lifestyle and gastrointestinal patterns, in an Indian cohort. Second, an *in-silico* comparative analysis was performed using curated dysbiosis records from the Disbiome database ^[17]^ for depression, anxiety, schizophrenia and ASD, to identify microbial taxa consistently elevated or reduced across these disorders and to place these taxa in phylogenetic context using NCBI taxonomy and the Interactive Tree of Life (iTOL) [19,20]. We hypothesized that disorders with overlapping symptomatology (depression, anxiety and schizophrenia) would show greater convergence in their dysbiosis signatures than the neurodevelopmentally distinct ASD.

The specific objectives of this study were to (i) assess public awareness of the gut microbiome’s role in mental health through a structured survey (ii) identify microbial taxa concordantly elevated or reduced in depression, anxiety, schizophrenia and ASD relative to healthy controls (iii) classify these taxa phylogenetically to reveal higher-order, phylum-level patterns underlying the observed dysbiosis.

## 2. Materials and Methods

### 2.1 Survey design and administration

An anonymous, structured questionnaire was designed using Google Forms to explore public understanding of the relationship between the gut microbiome and mental health. The form comprised three sections covering: (a) demographic details (age, gender, occupation and state of residence) (b) lifestyle and dietary information, including physical activity, consumption of gut-friendly/fermented foods, dietary restrictions, food preference under stress and self-rated mental well-being (0-10 scale) (c) medical history, including digestive symptoms during stress, family history of gastrointestinal or mental health conditions, frequency of doctor visits for gut-related concerns, prior mental health counselling and interest in learning more about the gut-brain connection. The survey was circulated through social media, e-mail and WhatsApp and administered in person, to reach participants of varying age, gender, occupation and geographic location. Participation was voluntary and responses were collected anonymously with no personal identification information analysis. Descriptive statistics (percentages) were used to summarise categorical responses.

### 2.2 Retrieval of microbial dysbiosis data

Disorder-specific microbial composition data for depression (major depressive disorder), anxiety, schizophrenia and ASD were retrieved from the Disbiome database (https://disbiome.ugent.be), a curated, literature-derived repository linking microbiome alterations to disease, maintained by Ghent University [17]. For each disorder, records were filtered to faecal samples only and each experimental entry was retrieved in JSON format containing the organism name, disease association and the qualitative outcome (elevated or reduced) relative to healthy controls. Each JSON record set was parsed and reorganized into two disorder-specific text files, one listing taxa reported as elevated and the other listing taxa reported as reduced, yielding eight lists in total (four disorders × two directions of change).

### 2.3 Comparative analysis of elevated and reduced taxa

The eight taxon lists were compared using the Multiple List Comparator tool (https://molbiotools.com/listcompare.php) [18], which computes pairwise and multi-way set intersections and generates a four-set Venn diagram. Separate comparisons were performed for the elevated-taxa lists and the reduced-taxa lists across the four disorders, yielding the number and identity of taxa shared between every combination of disorders. Taxa concordantly elevated or reduced in at least three of the four disorders were considered the core, shared dysbiosis signature and are reported in the main text; taxa shared between only two disorders are provided in the Supplementary Material (Tables S1 and S2).

### 2.4 Taxonomic classification and phylogenetic tree construction

The organism names comprising the shared elevated and reduced taxon sets were submitted to the NCBI Taxonomy tool (https://www.ncbi.nlm.nih.gov/taxonomy)[19] for classification and were exported in PHYLIP tree format using the NCBI Common Tree utility. The resulting files were visualized with the NCBI Phylogenetic Tree Viewer and exported in Newick format. Newick files were imported into the Interactive Tree of Life (iTOL v6) [20], where taxa were manually colour-annotated according to phylum-level assignment (Bacillota/Firmicutes, Actinomycetota/Actinomycetes, Bacteroidota/Bacteroidales, Pseudomonadota/Proteobacteria) to visualise the phylogenetic distribution of the shared dysbiotic taxa for the elevated and reduced sets separately.

### 2.5 Data handling and ethical considerations

No new wet-laboratory, animal, or human biological samples were generated for this study; all microbiome data were secondary, previously published records accessed through the Disbiome database. The survey component was conducted in accordance with standard ethical practice for anonymous, non-invasive online questionnaires, with informed voluntary participation and confidentiality of responses.

## 3. Results

### 3.1 Demographic profile and awareness of the gut-brain connection

The survey received 230 valid responses (Fig. 2). The majority of participants were aged between 20-30 years (53.9%), followed by the 30-40 (19.6%) and below-20 years (16.1%) age groups. Female participants predominated (70.4%) and nearly half of respondents (45.7%) were affiliated with the medical/health-sciences field, with a further 33% being students; this composition likely enhanced familiarity with the survey topic. Respondents were drawn predominantly from Rajasthan (87%), reflecting the location of the parent institution, with the remainder distributed across Delhi, Uttar Pradesh, Haryana, Karnataka and Maharashtra (Fig.2). Despite this geographic skew, 74.2% of respondents indicated awareness of a potential link between gut health and mental well-being, 12.2% were unaware and 13.5% were uncertain; unawareness was concentrated among students and respondents not affiliated with biology-related fields.

**Figure 1.**
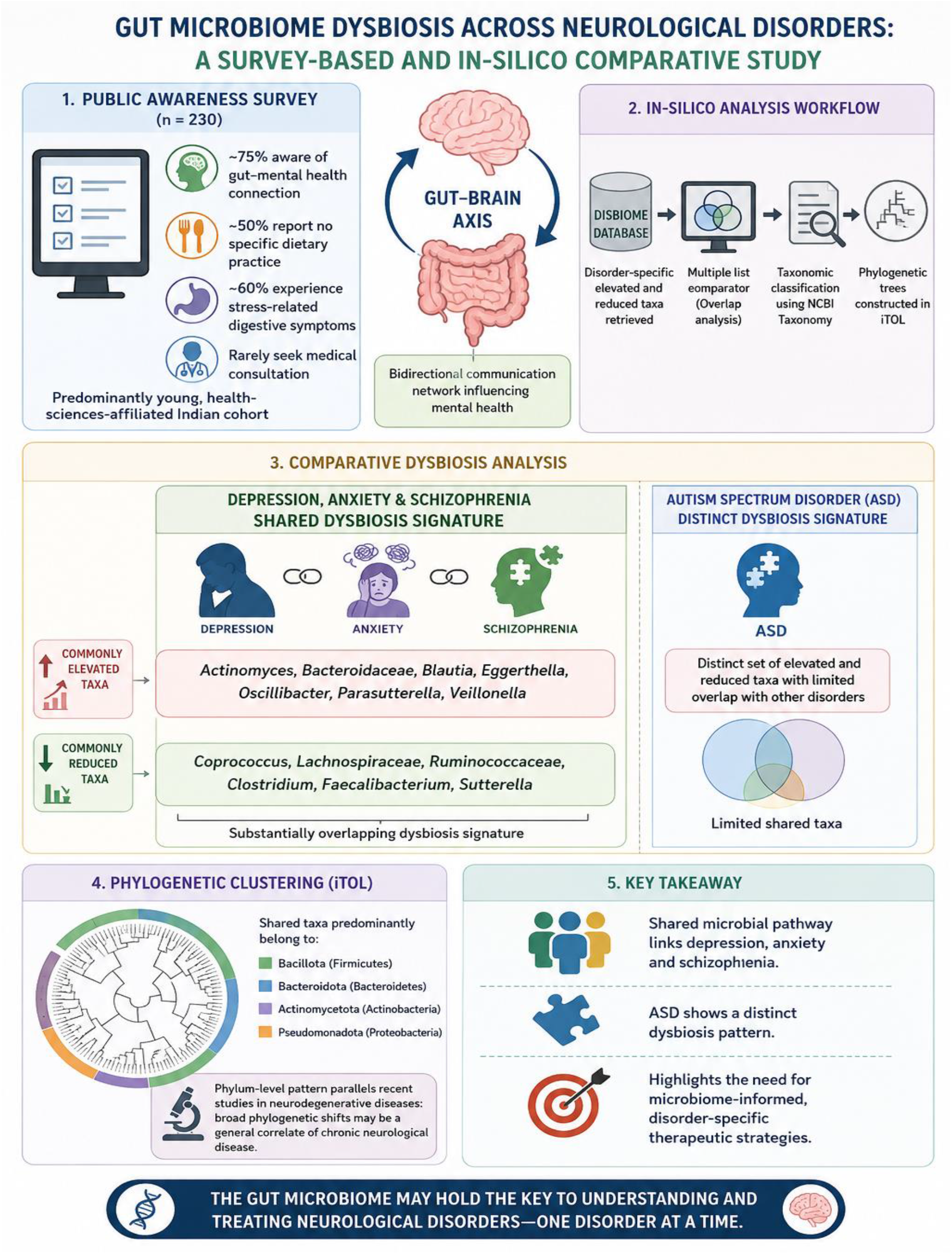
Graphical abstract summarizing the study workflow and key findings. The study combines a public-awareness survey (n = 230) with *in-silico* comparative analysis of gut microbiome dysbiosis in depression, anxiety, schizophrenia and autism spectrum disorder (ASD). Depression, anxiety and schizophrenia showed substantial overlap in microbial dysbiosis, whereas ASD displayed a distinct profile. Shared taxa predominantly clustered within *Bacillota, Bacteroidota, Actinomycetota* and *Pseudomonadota*, highlighting potential for disorder-specific, microbiome-informed therapeutic strategies.

**Figure 2.**
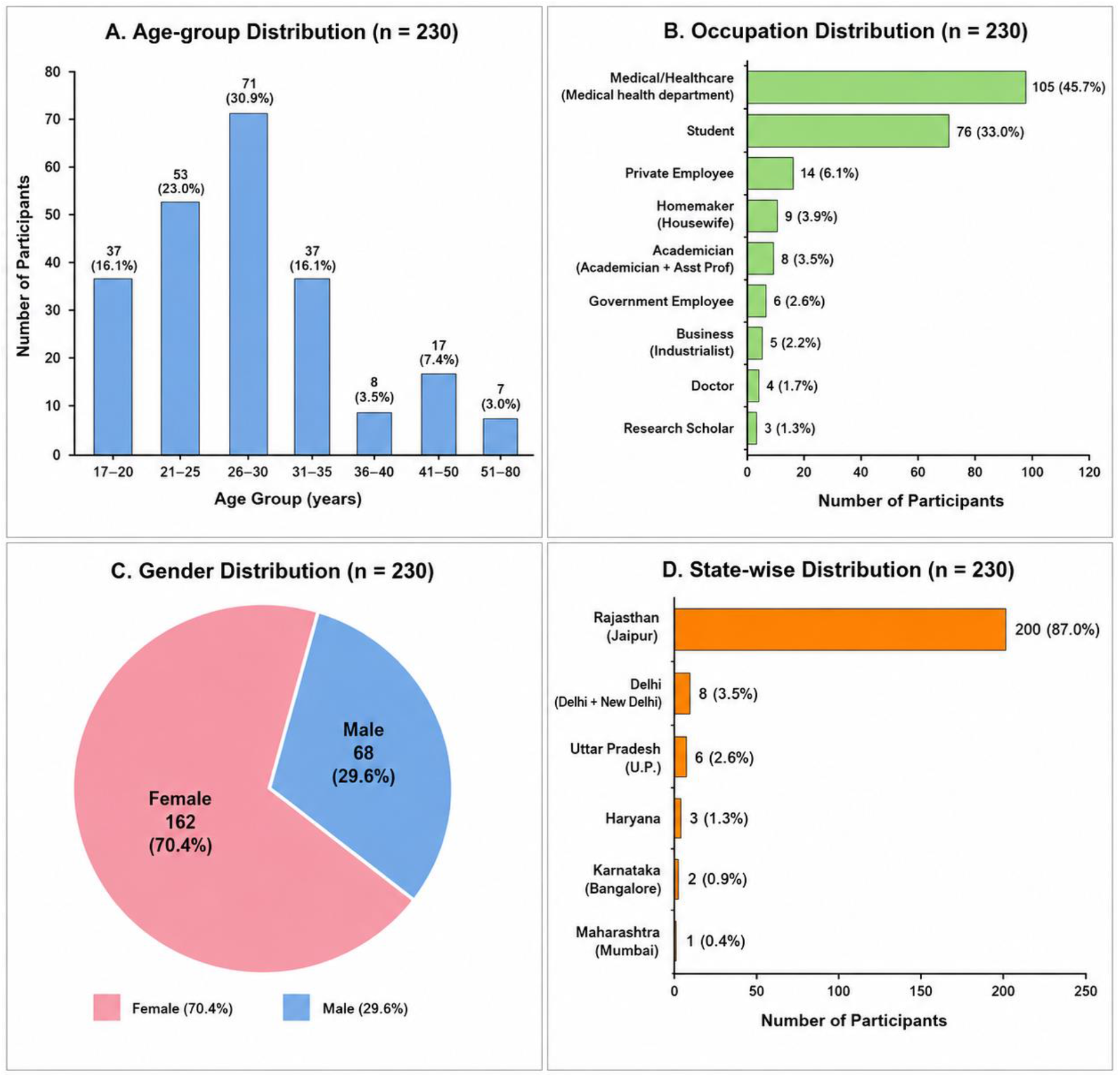
**Demographic profile of survey respondents (n = 230)**: age-group distribution, occupation, gender and state of residence.

### 3.2 Lifestyle, dietary and gastrointestinal correlates of stress

Approximately 74% of respondents reported regularly consuming gut-friendly or fermented foods such as yogurt and 65.2% engaged in regular physical activity, while just under half (49.1%) followed a specific dietary habit, typically involving avoidance of processed food and inclusion of fruits, vegetables and eggs. When queried about food preference during stress, more than half of respondents (52.6%) reported craving sweet foods, followed by spicy (24.8%), cheesy (15.7%) and salty (12.6%) foods, consistent with previously described stress-related shifts toward palatable, energy-dense food choices. Self-rated mental well-being scores were skewed toward the higher end of the 0-10 scale, with the modal rating of 8 (22.6% of respondents).

With respect to medical history, approximately 60% of respondents reported experiencing digestive symptoms, including acidity, bloating, indigestion, or irregular bowel movements, during periods of stress or anxiety, whereas participants engaging in regular exercise and consuming a nutritious diet reported comparatively fewer such symptoms. A family history of gastrointestinal or mental health conditions (including irritable bowel syndrome, generalised anxiety disorder and acidity) was reported by 14.3% of respondents. Despite the frequency of stress-related gastrointestinal symptoms, approximately 80% of respondents rarely or never sought medical consultation specifically for gut health, although a large majority (79.6%) expressed interest in learning more about the gut-brain connection, indicating a gap between perceived relevance and actual health-seeking behaviour.

### 3.3 Elevated microbial taxa shared across neurological disorders

Comparison of the elevated-taxa lists across depression (29 taxa), anxiety (36 taxa), schizophrenia (66 taxa) and ASD (107 taxa) identified thirteen taxa concordantly elevated in at least three of the four disorders (Table 1; Fig. 3A). Depression, anxiety and schizophrenia shared the largest overlap, including *Actinomyces, Bacteroidaceae, Blautia, Eggerthella, Oscillibacter, Parasutterella and Veillonella, whereas Ruminococcus* and *Shigella* were elevated across anxiety, schizophrenia and ASD. Pairwise intersection counts (Fig. 3A) confirmed that depression and anxiety shared the greatest number of elevated taxa (21), followed by schizophrenia-ASD (13) and anxiety-schizophrenia (11), while depression and ASD shared only four elevated taxa, the smallest pairwise overlap among all disorder combinations.

**Table 1.**
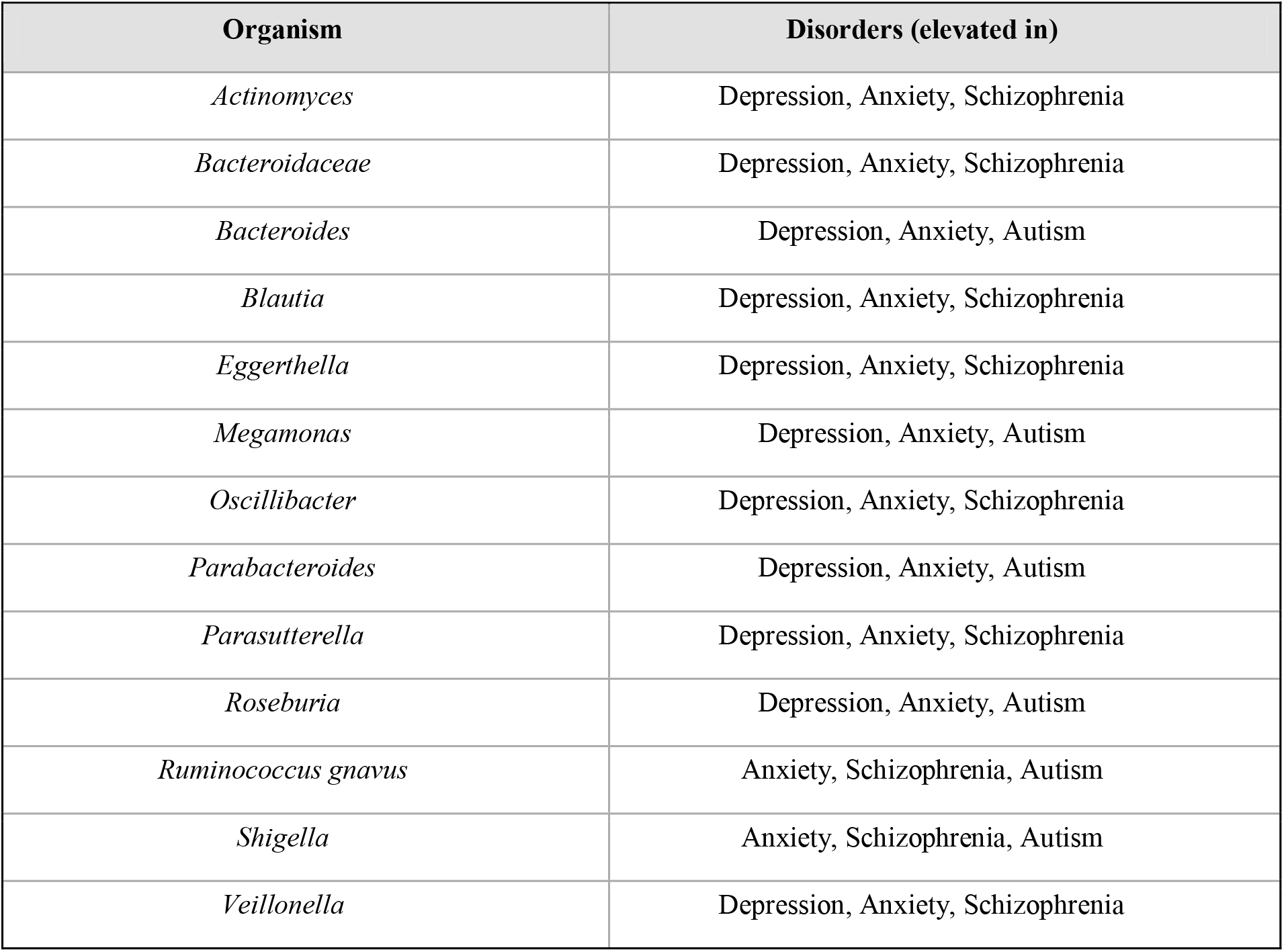
Microbial taxa concordantly elevated (relative to healthy controls) in at least three of the four neurological disorders examined.

| **Organism** | **Disorders (elevated in)** |
| --- | --- |
| *Actinomyces* | Depression, Anxiety, Schizophrenia |
| *Bacteroidaceae* | Depression, Anxiety, Schizophrenia |
| *Bacteroides* | Depression, Anxiety, Autism |
| *Blautia* | Depression, Anxiety, Schizophrenia |
| *Eggerthella* | Depression, Anxiety, Schizophrenia |
| *Megamonas* | Depression, Anxiety, Autism |
| *Oscillibacter* | Depression, Anxiety, Schizophrenia |
| *Parabacteroides* | Depression, Anxiety, Autism |
| *Parasutterella* | Depression, Anxiety, Schizophrenia |
| *Roseburia* | Depression, Anxiety, Autism |
| *Ruminococcus gnavus* | Anxiety, Schizophrenia, Autism |
| *Shigella* | Anxiety, Schizophrenia, Autism |
| *Veillonella* | Depression, Anxiety, Schizophrenia |

**Figure 3.**
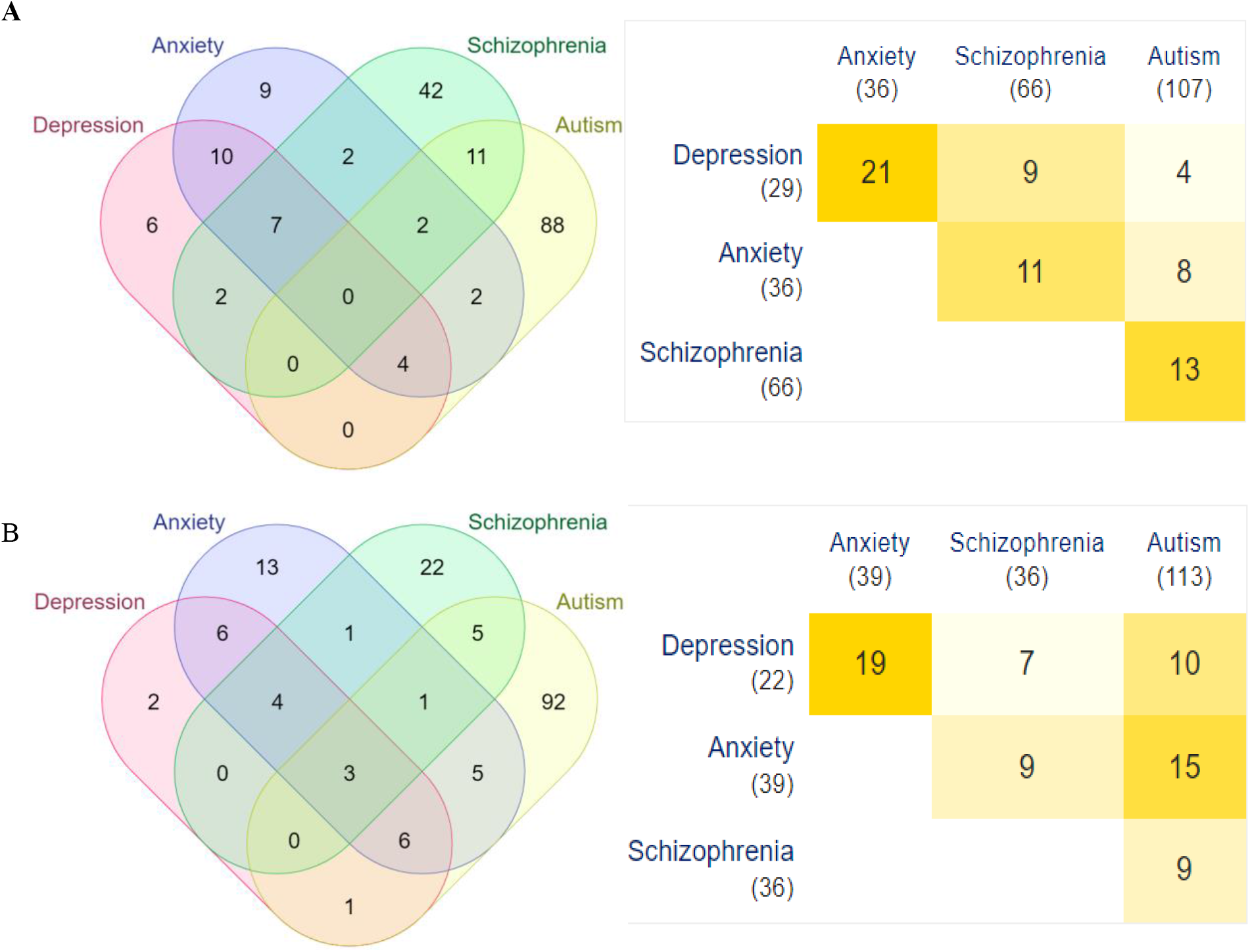
Four-way Venn diagrams and pairwise intersection matrices of microbial taxa. (A) elevated and (B) reduced across depression, anxiety, schizophrenia and autism spectrum disorder, generated using the Multiple List Comparator tool.

### 3.4 Reduced microbial taxa shared across neurological disorders

A parallel analysis of the reduced-taxa lists across depression (22), anxiety (39), schizophrenia (36) and ASD (113) identified fourteen taxa reduced in at least three disorders (Table 2; Fig. 3B), including three taxa (*Coprococcus, Lachnospiraceae* and *Ruminococcaceae*) reduced across all four disorders. As with the elevated set, depression and anxiety again shared the highest number of reduced taxa (19), whereas depression and schizophrenia shared the fewest (7). Anxiety and ASD shared 15 reduced taxa, the second-largest pairwise overlap observed, indicating a partial convergence between these two disorders at the level of depleted commensals despite their otherwise divergent dysbiosis profiles.

**Table 2.** Microbial taxa concordantly reduced (relative to healthy controls) in at least three of the four neurological disorders examined.

| **Organism** | **Disorders (reduced in)** |
| --- | --- |
| *Coprococcus* | Depression, Anxiety, Schizophrenia, Autism |
| *Lachnospiraceae* | Depression, Anxiety, Schizophrenia, Autism |
| *Ruminococcaceae* | Depression, Anxiety, Schizophrenia, Autism |
| *Bifidobacterium* | Depression, Anxiety, Autism |
| *Butyricicoccus* | Anxiety, Schizophrenia, Autism |
| *Clostridium* | Depression, Anxiety, Schizophrenia |
| *Dialister* | Depression, Anxiety, Autism |
| *Enterobacteriaceae* | Depression, Anxiety, Schizophrenia |
| *Faecalibacterium* | Depression, Anxiety, Schizophrenia |
| *Lactobacillus* | Depression, Anxiety, Autism |
| *Phascolarctobacterium* | Depression, Anxiety, Autism |
| *Prevotella* | Depression, Anxiety, Autism |
| *Roseburia* | Depression, Anxiety, Autism |
| *Sutterella* | Depression, Anxiety, Schizophrenia |

### 3.5 Phylogenetic organization of the shared dysbiotic taxa

Taxonomic classification and phylogenetic reconstruction of the shared elevated taxa (Fig. 4A) showed that the majority belonged to the phylum Bacillota (Firmicutes), including genera such as *Blautia, Oscillibacter, Roseburia, Veillonella* and *Megasphaera*. Additional representation came from Actinomycetota (Actinomycetes; e.g., *Eggerthella, Coriobacteriaceae, Bifidobacteriaceae*), Bacteroidota (Bacteroidales; e.g., *Alistipes, Parabacteroides, Bacteroides fragilis*) and Pseudomonadota (Proteobacteria; e.g., *Parasutterella, Burkholderia, Shigella*).

**Figure 4.**
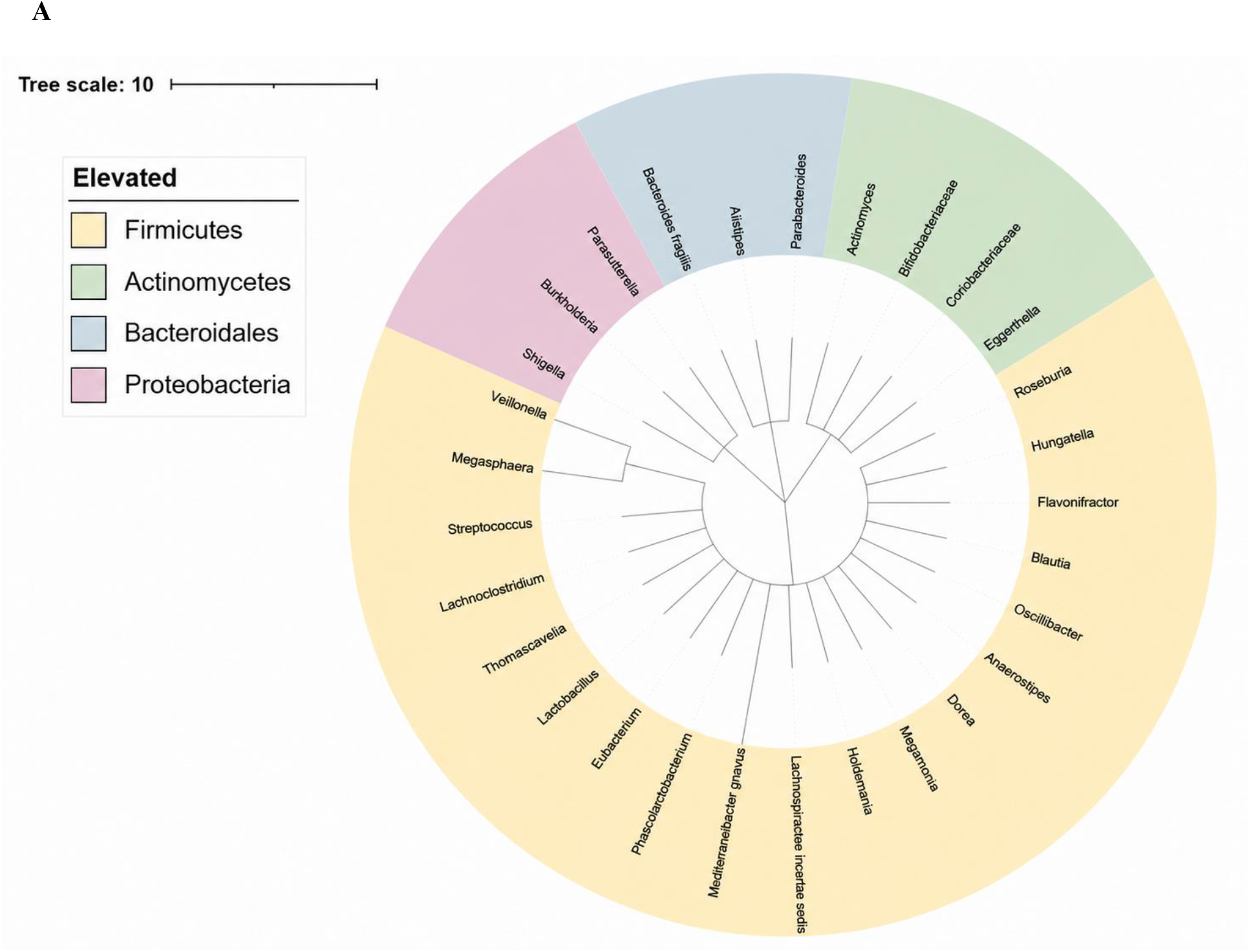
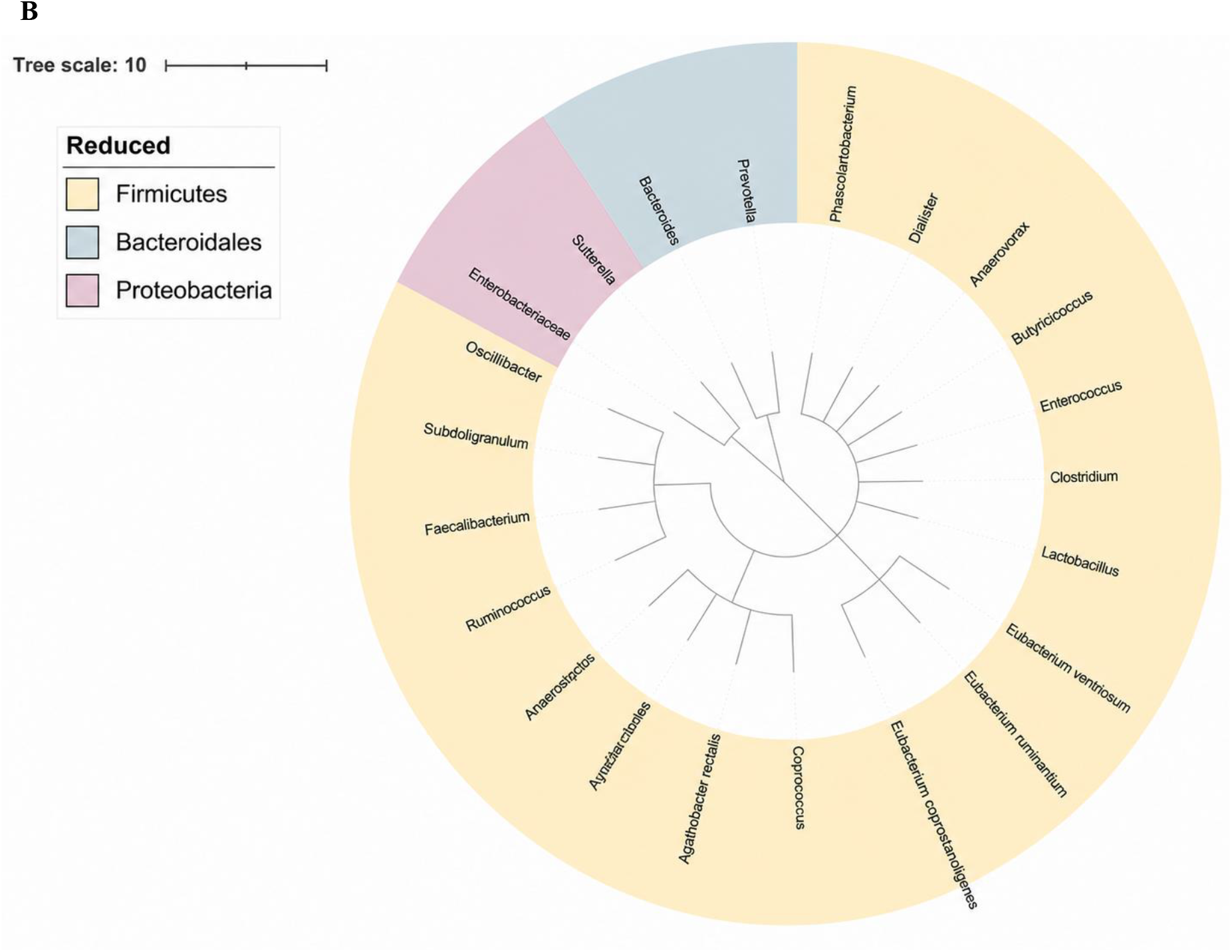
Phylogenetic distribution, coloured by phylum of taxa shared across at least three neurological disorders: (A) elevated taxa (B) reduced taxa. Trees were constructed via NCBI Taxonomy/Common Tree and visualised in iTOL.

The shared reduced taxa (Fig. 4B) also clustered predominantly within Bacillota, including genera such as *Faecalibacterium, Clostridium, Lactobacillus* and *Coprococcus*, with smaller contributions from Bacteroidota (*Prevotella, Bacteroides*) and Pseudomonadota (*Sutterella, Enterobacteriaceae*). Notably, no Actinomycetota members were present among the reduced taxa.

Overall, both the elevated and reduced dysbiotic signatures were dominated by phylum Bacillota, consistent with its status as the most abundant phylum in the healthy human gut microbiome and therefore the phylum most likely to exhibit detectable bidirectional shifts during disease states.

## 4. Discussion

This study combined primary survey data with a secondary, in-silico comparative analysis to examine both public perception and molecular evidence for the gut-brain relationship across four neurological disorders. Three-quarters of respondents were already aware of a potential gut-mental health link, considerably higher than might be expected from general population sampling, which likely reflects the health-sciences-weighted composition of the surveyed cohort rather than broader public awareness in India. Even within this relatively informed group, however, a clear gap was evident between recognising the gut–brain connection and acting on it: most respondents rarely sought medical attention for stress-related digestive symptoms despite their high reported frequency (∼60%), suggesting that gastrointestinal symptoms continue to be perceived as a secondary consequence of stress rather than a modifiable contributor to it.

This behavioural gap raises a more fundamental question: is there a plausible biological basis for treating these stress-linked digestive symptoms as clinically meaningful rather than incidental and does this basis differ across neuropsychiatric and neurodevelopmental conditions? The *in-silico* comparative analysis was designed to address this question directly by asking whether the microbial changes reported in depression, anxiety, schizophrenia, and ASD converge on a shared, biologically coherent signature, or whether they are too disorder-specific to support any general “gut-health” recommendation of the kind the survey suggests respondents are not currently receiving or acting on. The results indicate that both are partly true. The comparative dysbiosis analysis revealed a substantially overlapping microbial signature among depression, anxiety, and schizophrenia, both in the taxa elevated and in those reduced relative to healthy controls, whereas ASD exhibited a distinct profile with markedly less overlap This pattern is reflected in the shared elevation of Bacteroidaceae, *Blautia, Eggerthella, Oscillibacter, Parasutterella* and *Veillonella*, together with the shared depletion of *Coprococcus*, Lachnospiraceae, Ruminococcaceae, *Faecalibacterium, Clostridium* and *Sutterella* across these three disorders. Several of these taxa have plausible mechanistic links to psychiatric symptoms. *Faecalibacterium* and *Coprococcus* are major butyrate producers; their consistent depletion across depression, anxiety and schizophrenia is compatible with reduced short-chain-fatty-acid-mediated anti-inflammatory and neurotrophic signalling, including diminished support for Brain-Derived Neurotrophic Factor (BDNF) expression [10]. Notably, this same loss of butyrate-producing capacity provides a concrete mechanistic candidate for the digestive symptoms reported by the majority of survey respondents during stress: reduced butyrate output weakens gut barrier integrity and colonic health, offering a biologically grounded reason why the “stress causes stomach upset” narrative volunteered by several respondents (Supplementary Text S1) may in fact run in both directions. Conversely, elevation of Proteobacteria-associated genera such as *Parasutterella* and *Shigella*-like organisms may indicate increased gut barrier permeability (“leaky gut”) and low-grade systemic inflammation, both of which have been proposed as mediators of neuroinflammation in depression, anxiety and schizophrenia [7,10,11,13]. The concordant reduction of Lachnospiraceae and Ruminococcaceae, both major contributors to short-chain fatty acid production and mucosal integrity, across all four disorders, including ASD, suggests that loss of these butyrate-producing families may be a fairly general sign of neurological dysfunction, even when the rest of the dysbiosis pattern differs between disorders.

The comparatively limited overlap between ASD and the other three disorders, both for elevated and reduced taxa, mirrors its distinct pathophysiological origin: ASD is fundamentally a neurodevelopmental condition with onset in early childhood, during a critical window in which the gut microbiome itself is still being established, whereas depression, anxiety, and schizophrenia typically manifest later and may share more overlapping downstream neuroimmune and neuroendocrine mechanisms in adulthood [9,12]. This distinction reinforces the conclusion, drawn previously for individual disorders [10,11,12,13], that microbiome-informed interventions are unlikely to be interchangeable across all neurological conditions and instead require disorder-specific validation. It also refines the interpretation of the survey’s awareness–action gap: generic advice to “look after your gut” for mental well-being is not equally well-supported across conditions, and closing this gap meaningfully will require condition-specific public-health messaging rather than a single undifferentiated message linking gut and mental health.

At the phylum level, both the elevated and reduced taxon sets were dominated by Bacillota (Firmicutes), together with Bacteroidota (Bacteroidetes), which under normal physiological conditions constitute the two most abundant phyla in the healthy human gut, typically accounting for the large majority of the total bacterial community [1,5]. This baseline dominance likely explains, at least in part, why shifts within these two phyla are so consistently detected across disease states: any perturbation of the gut ecosystem is statistically more likely to register as a change among its most abundant residents. Nonetheless, the consistent bidirectional involvement of specific Bacillota genera, for example, simultaneous elevation of some genera (Blautia, Oscillibacter) and depletion of others (Faecalibacterium, Clostridium), indicates that the observed dysbiosis reflects a genuine functional reorganisation within this phylum rather than a simple net expansion or loss. This is consistent with the broader understanding that the gut microbiome’s contribution to host physiology, including nutrient metabolism, short-chain fatty acid production, and immune modulation, depends on the relative balance among functionally distinct taxa within a phylum rather than on phylum-level abundance alone [1,6].

Several limitations should be considered. The survey sample was geographically concentrated in Rajasthan and disproportionately represented health-sciences-affiliated and female respondents, limiting generalisability to the broader Indian population; a more demographically balanced and larger sample would strengthen future survey-based assessments. The microbiome comparison relied on secondary, literature-curated qualitative outcome data (elevated/reduced) from Disbiome rather than raw sequencing data, precluding effect-size or statistical-significance-weighted comparison and introducing potential heterogeneity arising from differences in detection method, sample type, and control definitions across the underlying primary studies. Because the survey and the microbiome data were drawn from independent sources rather than the same individuals, this study cannot directly test whether the respondents’ own symptom patterns correspond to the dysbiosis signatures described here; this linkage remains inferential and should be confirmed in future work that pairs symptom or diagnostic data with microbiome sampling in the same participants. The in-silico nature of the comparison also cannot establish causality between microbial dysbiosis and disease; longitudinal and mechanistic studies, ideally paired with individual-level survey data such as those collected here, would be required to determine whether dysbiosis precedes, accompanies, or results from these neurological conditions. Finally, the present analysis was restricted to four disorders and did not include comparator dysbiosis profiles from healthy individuals or non-neurological disease states, which would help establish the specificity of the identified taxa to neurological, as opposed to general inflammatory, disease processes.

Despite these limitations, this comparative approach demonstrates the utility of integrating publicly available curated microbiome databases with taxonomic and phylogenetic tools to identify candidate, cross-disorder microbial signatures efficiently and without new sequencing, and of pairing this molecular picture with real-world survey data on awareness and health behaviour. Future work should prioritise quantitative meta-analysis of raw abundance data were available, functional (metagenomic or metabolomic) characterisation of the shared taxa identified here, particularly their genomic capacity for short-chain fatty acid and neurotransmitter-precursor biosynthesis, and intervention studies targeting the shared depleted butyrate producers (Faecalibacterium, Coprococcus) as a rational, disorder-spanning therapeutic strategy for depression, anxiety, and schizophrenia. Such studies, conducted in cohorts where survey-reported symptoms and microbiome composition are measured in the same individuals, would also help translate the awareness gap identified here into targeted, biologically justified public-health guidance.

## 5. Conclusion

This study integrated a public-awareness survey with a comparative *in-silico* analysis of gut microbial dysbiosis across four neurological disorders: depression, anxiety, schizophrenia and autism spectrum disorder (ASD). Although most of the surveyed population recognized the existence of a gut–brain connection, this awareness rarely translated into preventive health-seeking behaviour, pointing to a gap between knowledge and action that future public-health messaging could usefully address.

At the biological level, the findings carry several meaningful implications. Depression, anxiety and schizophrenia shared a substantially convergent pattern of dysbiosis, marked by loss of key butyrate-producing taxa such as *Faecalibacterium* and *Coprococcus*. Because butyrate and related short-chain fatty acids help maintain gut barrier integrity, dampen systemic inflammation and support neurotrophic signalling in the brain, their consistent depletion across these three disorders points to a shared biological pathway, reduced short-chain-fatty-acid output, that could plausibly contribute to the low-grade inflammation and impaired neuroplasticity reported in each condition individually. At the same time, the elevation of Proteobacteria-associated genera such as *Parasutterella* across these disorders is consistent with increased gut permeability and a pro-inflammatory microbial environment, reinforcing inflammation as a candidate mechanistic link between the gut and the brain in these conditions.

ASD, by contrast, showed a largely distinct microbial signature, in keeping with its origin as a neurodevelopmental condition that emerges while the gut microbiome itself is still being established in early life, rather than a disorder arising from later-life neuroimmune or neuroendocrine changes. This biological distinction is clinically important: it suggests that microbiome-targeted interventions such as probiotics, prebiotics or dietary modification are unlikely to work identically across all four disorders, and that treatments aimed at restoring depleted butyrate producers may hold shared therapeutic value for depression, anxiety and schizophrenia specifically, while ASD will likely require a separately tailored approach.

More broadly, this work reinforces the biological plausibility of the gut-brain axis as a contributor to neuropsychiatric and neurodevelopmental disease, operating through interconnected short-chain-fatty-acid, neurotransmitter and immune-signalling pathways. Larger, demographically representative surveys and quantitative, sequencing-based studies of these same taxa, particularly their genomic capacity to produce short-chain fatty acids and neurotransmitter precursors, are needed to move from this associative, *in-silico* picture toward mechanistic understanding and, ultimately, toward microbiome-informed diagnostic and therapeutic strategies for these disorders.

## Acknowledgements

The authors sincerely thank all survey participants for their voluntary contribution.

## Data Availability Statement

The microbial dysbiosis data analysed in this study are publicly available from the Disbiome database (https://disbiome.ugent.be). Survey data supporting the findings of this study are available from the corresponding author upon reasonable request.

