## Supplementary Material for "Deciphering the Gut–Brain Dialogue: A Survey-Based and *In-Silico* Comparative Analysis of Gut Microbial Dysbiosis in Common Neurological Disorders"

**Supplementary Table S1**

Microbial taxa reported as elevated (relative to healthy controls) in exactly two of the four neurological disorders examined (depression, anxiety, schizophrenia, autism spectrum disorder). Taxa elevated in three or four disorders are reported in Table 1 of the main text.

| **Organism** | **Disorders (elevated in)** |
| --- | --- |
| *Akkermansia muciniphila* | Schizophrenia, Autism |
| *Alistipes* | Depression, Anxiety |
| *Anaerostipes* | Depression, Anxiety |
| *Bacteroidales* | Depression, Anxiety |
| *Bacteroides fragilis* | Schizophrenia, Autism |
| *Bacteroidetes* | Depression, Anxiety |
| *Bifidobacteriaceae* | Depression, Anxiety |
| *Bilophila* | Schizophrenia, Autism |
| *Burkholderia* | Anxiety, Autism |
| *Clostridium cluster XVIII* | Depression, Anxiety |
| *Clostridium innocuum* | Schizophrenia, Autism |
| *Clostridium symbiosum* | Schizophrenia, Autism |
| *Coriobacteriaceae* | Depression, Schizophrenia |
| *Dorea* | Schizophrenia, Autism |
| *Enterobacteriaceae* | Anxiety, Autism |
| *Eubacterium siraeum* | Schizophrenia, Autism |
| *Eubacterium* | Depression, Anxiety |
| *Flavonifractor* | Depression, Schizophrenia |
| *Holdemania* | Schizophrenia, Autism |
| *Hungatella* | Anxiety, Schizophrenia |
| *Lachnospiraceae incertae sedis* | Depression, Anxiety |
| *Lactobacillus* | Schizophrenia, Autism |
| *Megasphaera* | Schizophrenia, Autism |
| *Methanobrevibacter* | Anxiety, Schizophrenia |
| *Phascolarctobacterium* | Depression, Anxiety |
| *Streptococcus* | Depression, Anxiety |
| *Veillonellaceae* | Schizophrenia, Autism |

**Supplementary Table S2**

Microbial taxa reported as reduced (relative to healthy controls) in exactly two of the four neurological disorders examined. Taxa reduced in three or four disorders are reported in Table 2 of the main text.

| **Organism** | **Disorders (reduced in)** |
| --- | --- |
| *Agathobacter* | Anxiety, Schizophrenia |
| *Anaerostipes* | Anxiety, Autism |
| *Anaerovorax* | Depression, Anxiety |
| *Bacteroides* | Depression, Anxiety |
| *Bacteroidetes* | Depression, Autism |
| *Clostridium cluster XVIII* | Depression, Anxiety |
| *Clostridium group XI* | Depression, Anxiety |
| *Enterococcus* | Schizophrenia, Autism |
| *Eubacterium coprostanoligenes* | Anxiety, Autism |
| *Eubacterium rectale* | Anxiety, Autism |
| *Eubacterium ruminantium* | Anxiety, Autism |
| *Eubacterium ventriosum* | Schizophrenia, Autism |
| *Fusicatenibacter* | Schizophrenia, Autism |
| *Fusobacterium* | Schizophrenia, Autism |
| *Oscillibacter* | Depression, Anxiety |
| *Roseburia faecis* | Schizophrenia, Autism |
| *Ruminococcus* | Depression, Anxiety |
| *Subdoligranulum* | Anxiety, Autism |

**Survey Data - Supplementary Figures**

The following figures present the complete survey results (n = 230 valid responses) beyond the demographic profile shown in Figure 2 of the main text. All charts are horizontal bar charts with response categories on the vertical axis and the percentage of respondents on the horizontal axis; the horizontal axis is individually scaled just above each chart's highest value so that bars fill the plot area meaningfully, and no numeric labels are printed on the bars — values are read directly from the percentage axis.

**Supplementary Figure S1**

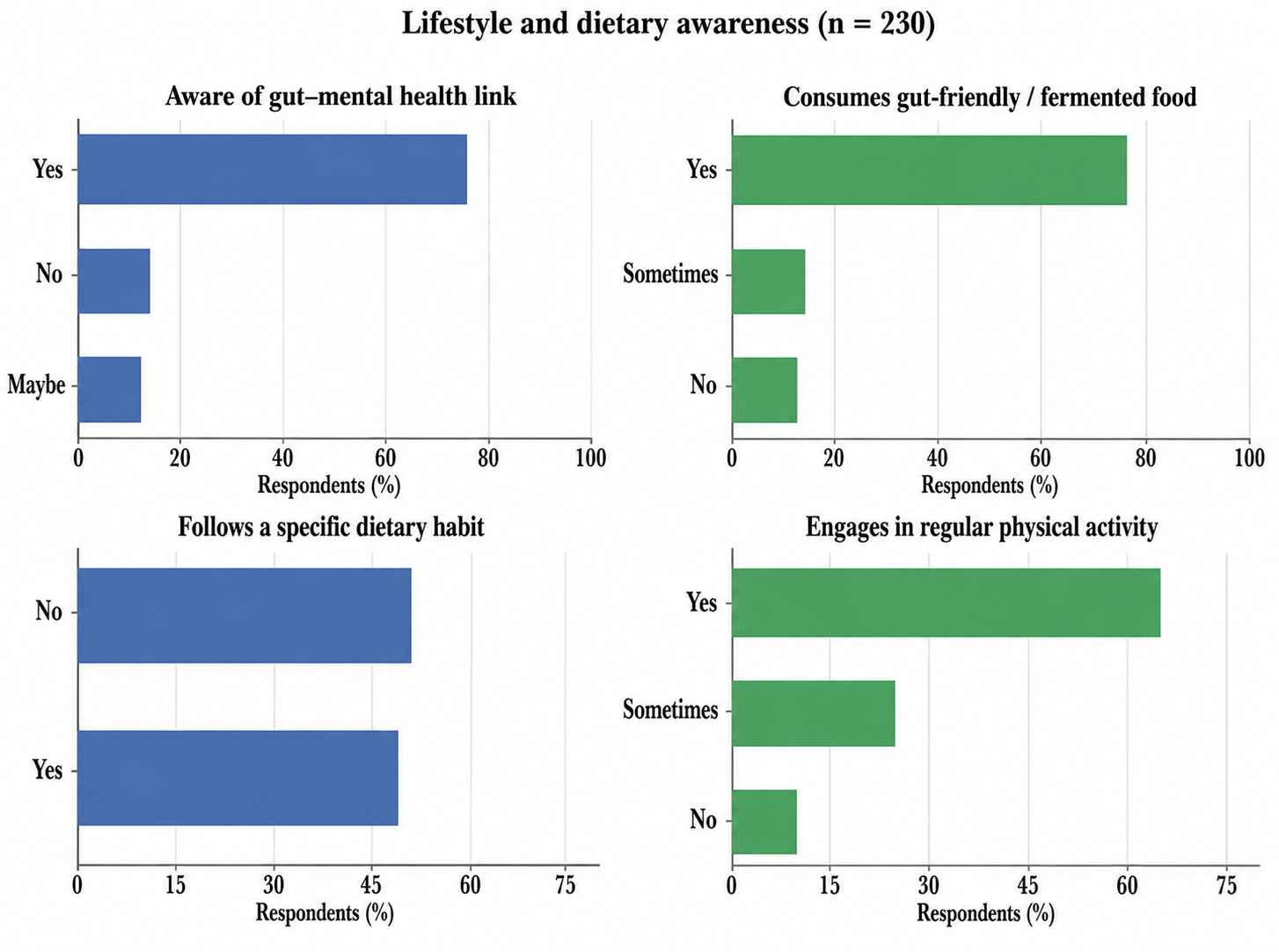

**Figure S1**. Awareness of the gut–mental health link and lifestyle habits potentially relevant to gut health (n = 230).

**Observations.** Nearly three-quarters of respondents (73.9%) reported awareness that gut health may affect mental well-being, with a further 12.2% unsure (‘Maybe’) and only 13.9% unaware. Reported gut-friendly/fermented food consumption is similarly high (73.5% ‘Yes’), and 65.2% report regular physical activity. Only about half of respondents (49.1%) follow any specific dietary habit or restriction intended to support gut health, indicating that awareness of the gut–brain link does not automatically translate into a deliberate, described dietary strategy for around half the sample.

**Supplementary Figure S2**

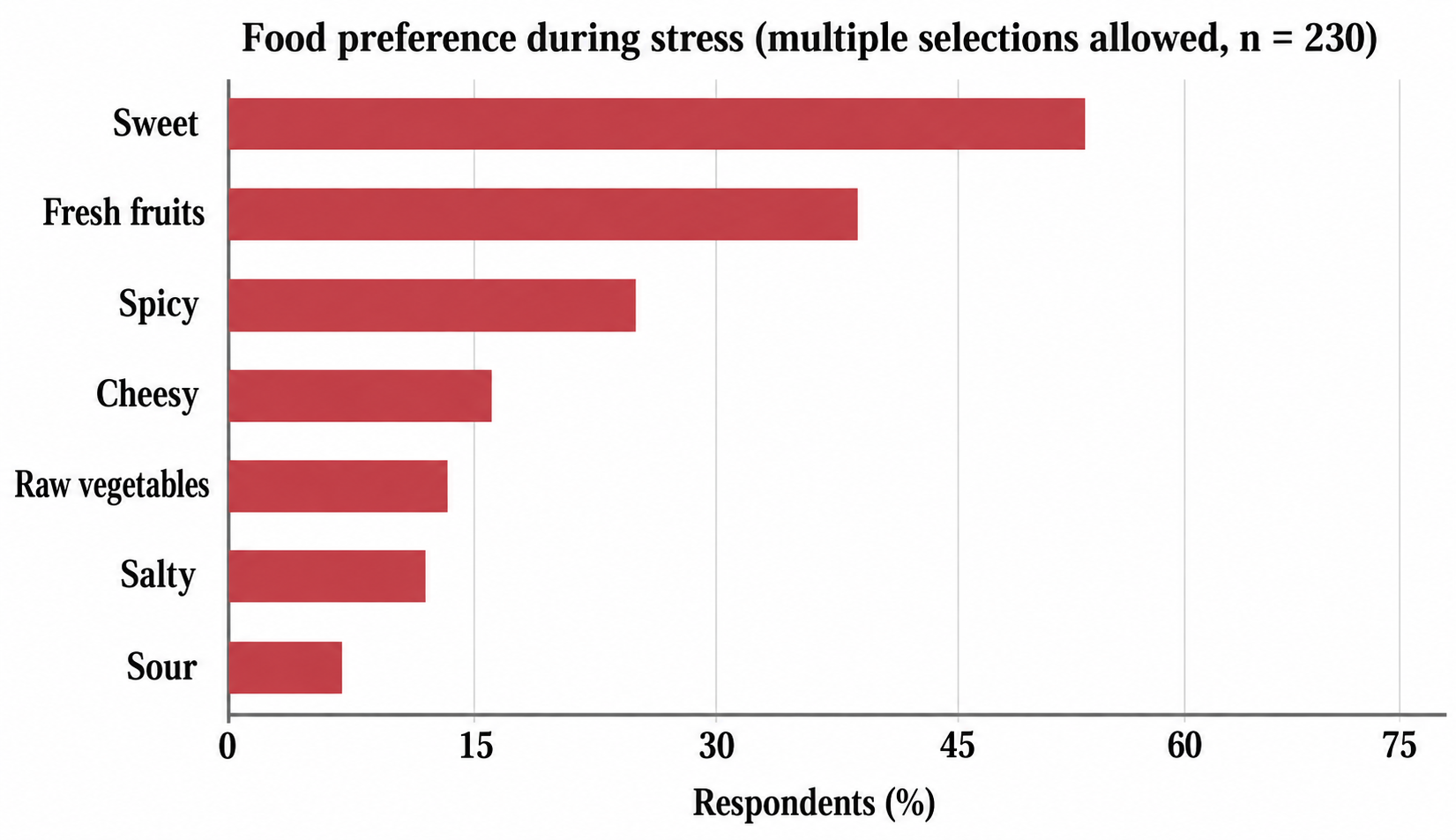

**Figure S2.** Self-reported food preference during stress (n = 230; respondents could select more than one category, so percentages do not sum to 100%).

**Observations.** Sweet food is the dominant stress-related craving, selected by 52.6% of respondents, more than double the next most common choice (fresh fruit, 37.8%). Spicy (24.8%), cheesy (15.7%), raw vegetables (13.5%), salty (12.6%), and sour (7.0%) foods were selected less often, suggesting that both highly palatable, energy-dense choices (sweet) and fibre-rich choices (fresh fruit) coexist as stress-eating tendencies in this sample.

**Supplementary Figure S3**

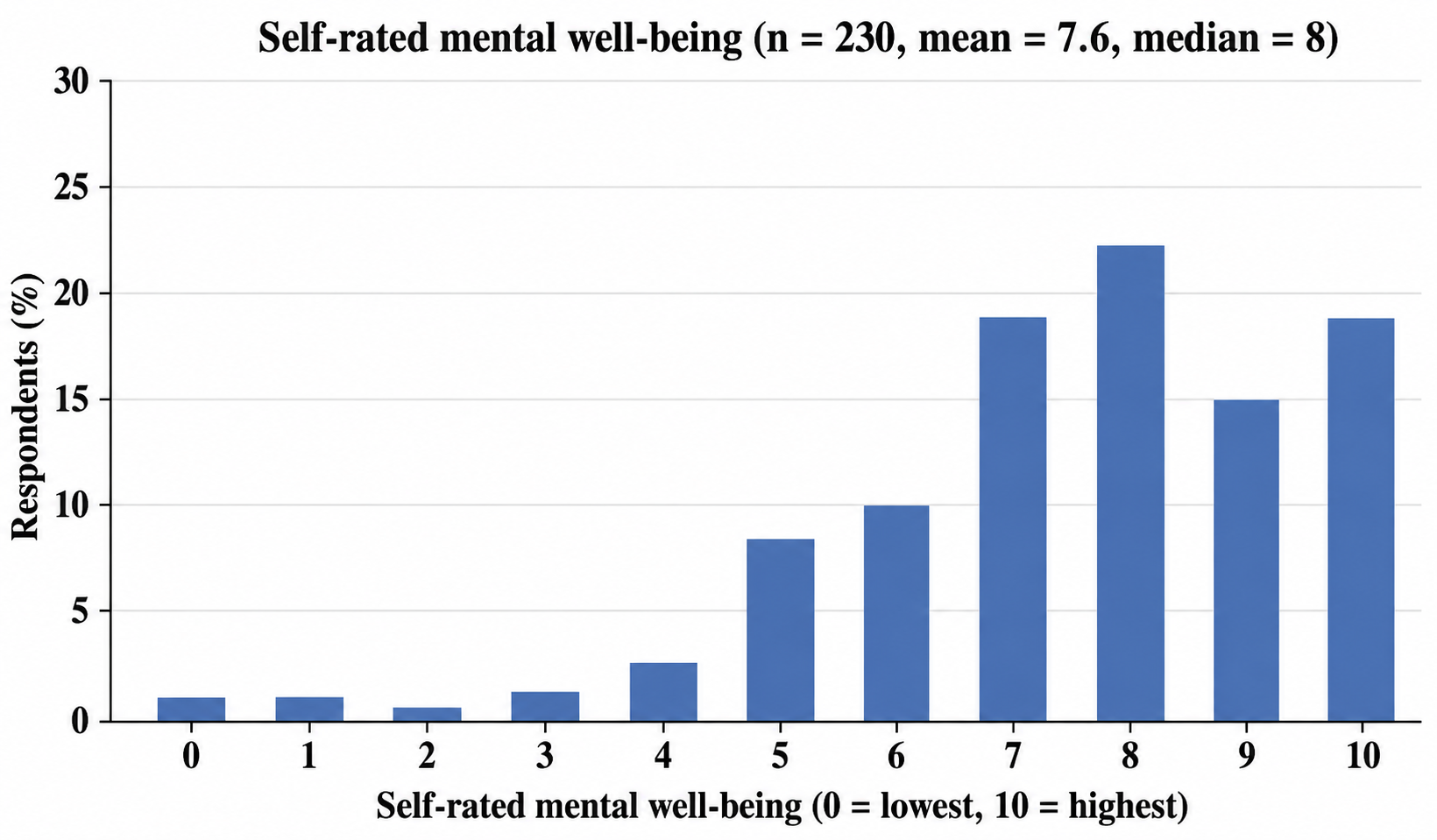

**Figure S3.** Distribution of self-rated overall mental well-being on a 0 (lowest) to 10 (highest) scale (n = 230; mean = 7.56, median = 8).

**Observations.** Self-rated well-being is skewed toward the higher end of the scale: scores of 7–10 account for 75.6% of all responses, with 8 the single most common rating (22.6%). Low scores are rare — ratings of 0–2 together account for only 2.2% of respondents — consistent with a generally healthy, non-clinical convenience sample rather than one enriched for diagnosed mental health conditions.

**Supplementary Figure S4**

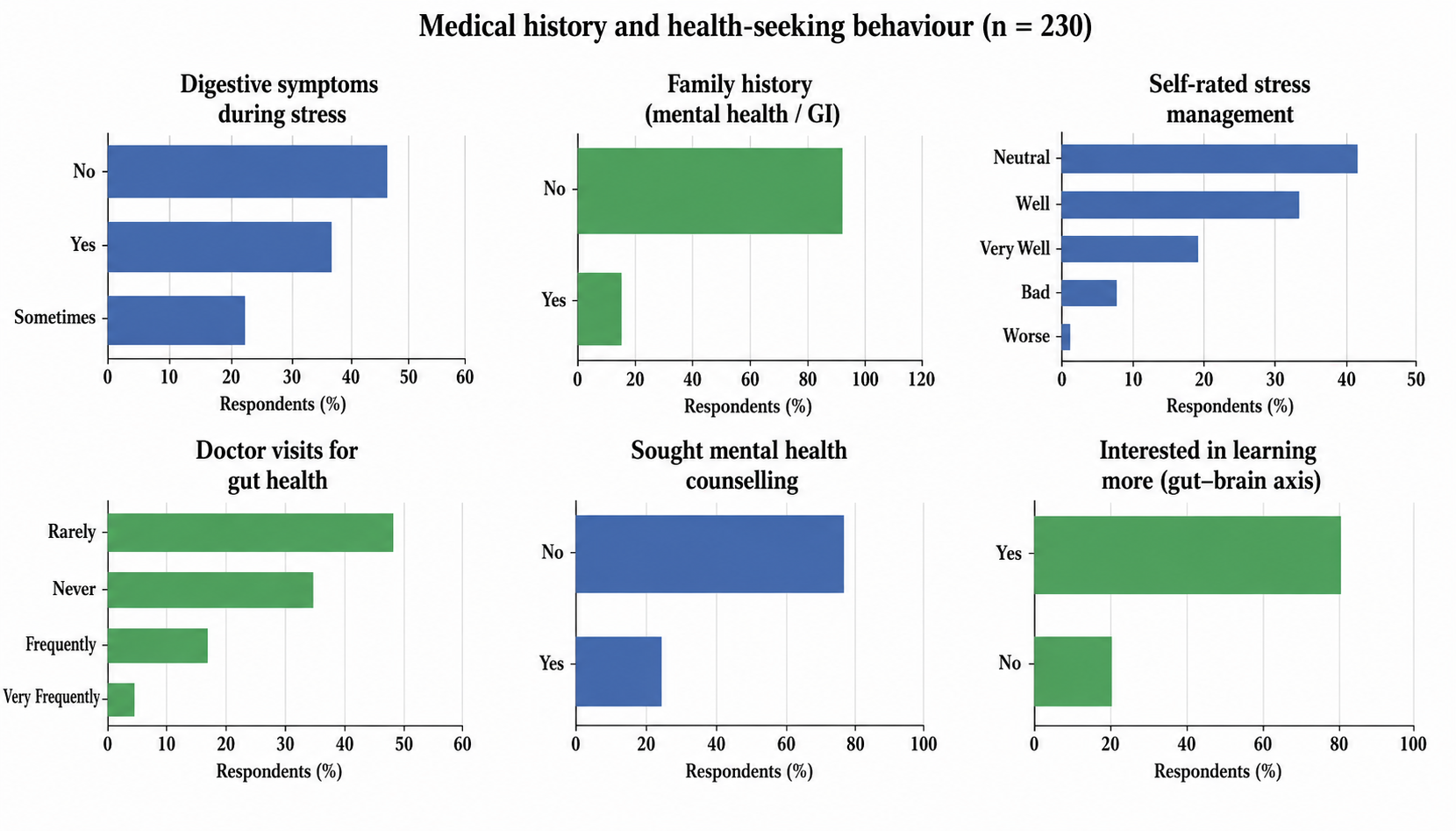

**Figure S4**. Medical history and health-seeking behaviour (n = 230): digestive symptoms during stress, family history, self-rated stress management, doctor visits for gut health, prior counselling, and interest in learning more.

**Observations.** 57.4% of respondents experience digestive symptoms at least sometimes during stress (36.1% ‘Yes’ + 21.3% ‘Sometimes’), yet 80.0% rarely or never see a doctor specifically for gut health, and only 23.9% have ever sought professional counselling for a mental health concern. A family history of a mental health or gastrointestinal condition was reported by 14.3% of respondents. Despite low rates of professional consultation, interest in learning more about the gut–brain connection is high (79.6%), pointing to an awareness–action gap between symptom prevalence/interest and formal health-seeking behaviour.

**Supplementary Text S1: Illustrative Free-Text Survey Responses**

Two open-ended questions allowed respondents to elaborate in their own words. Representative responses are reproduced below; full verbatim responses are available from the corresponding author upon request.

**S1.1 “Would you like to specify your dietary habit?” (30 of 230 respondents answered)**

*“I prefer fruits like apple and banana, including yogurt, green vegetables and kefir grains in my diet.”*

*“Always take curd, salad in your lunch.”*

*“Including curd or buttermilk once daily.”*

*“No milk or soft drink after having curd.”*

*“People having certain digestive disorders are prone to certain mental issues such as depression and anxiety, so there must be a balancing of bacteria … there must be proper dietary rules for every citizen.”*

**S1.2 “Family history of mental health or gastrointestinal conditions?” (21 of 230 respondents specified)**

*“My father is suffering with Parkinson's.”*

*“My mother has been facing gastric problems since the last few years, and whenever she is stressed or overthinks, her stomach gets upset.”*

*“Depression and IBS.”*

*“GAD (generalised anxiety disorder).”*

*“Stress and acidity is very common [in my family].”*

*“Overthinking.”*
